# Early visual experience elicits cellular and functional plasticity in the retina and alters behaviour

**DOI:** 10.1101/2025.04.29.651180

**Authors:** Phoebe Reynolds, Davide Marchi, Yan To Ling, Katja Slangewal, Max Capelle, Zhaklin Chalakova, Armin Bahl, Robert Hindges

## Abstract

Our interaction with the surrounding environment shapes how our brain processes sensory information and drives adaptive behaviour. This plasticity allows the brain to rewire in response to specific sensory experiences. For instance, early manipulation of visual inputs profoundly impacts brain plasticity, which is crucial for functions like size perception, object recognition, and visuospatial processing. While neuronal plasticity has been detected in visual target structures such as the colliculus, thalamus, and cortex, it remains unclear if the retina, the primary sensory organ, undergoes significant plasticity. Here, we show that the zebrafish retina demonstrates pronounced plastic transformations in response to alterations of the visual environment during development, which ultimately modifies the detection of oriented visual stimuli. We demonstrate that orientation-selective amacrine cells undergo profound morphological changes in animals exposed to distinct visual environments during development. We further find that the functional orientation-selective output from the retina is altered in a manner consistent with the visual environment in which the animals are raised and that these changes are persistent. Finally, animals tested in a virtual reality system show that early exposure to different visual environments changes their innate preference for specifically oriented patterns. Our findings unveil a unique developmental form of sensory organ plasticity with continuing structural and functional consequences.

## Introduction

Our behaviour relies on the appropriate processing of sensory information. Visual perception is dependent on the intricate circuitry of the visual system to navigate and interact with our environment accordingly^1–3^, which is established through a combination of molecular, cellular, and activity-dependent processes during development^4^. A crucial component is the ability to adapt sensory processing to changing environmental conditions, to possibly gain an evolutionary advantage^5^. This interplay between experience and circuitry is evident across different sensory modalities^6–8^ and, within the visual circuit, is known to profoundly impact upon higher-order visual processing in humans^9–11^. Although it is widely recognised that investigating these dynamics is crucial for understanding the mechanisms that drive visual adaptive responses^12–14^, the full extent of plasticity across the visual system is not clear.

Research into visual plasticity has focused largely on the malleability of cortical structures, and it is largely agreed that sensory experiences significantly influence the functional organisation of visual circuits during maturation^15–20^. However, the potential for plasticity at the level of the sensory organs themselves, particularly the retina, remains largely unexplored, as the retina has been considered to possess a rigid structure that provides a stable foundation for changes occurring elsewhere^21^. Recent studies have suggested a disconnect between activity-dependent plasticity and behavioural outputs^22,23^, with the greatest extent of dynamic changes within the retina primarily considering the role of spontaneous retinal wave activity^24,25^ and altered sensory sensitivity through visual deprivation^16,26–28^. However, given the retina’s pivotal role in extracting salient visual information^29–33^, this raises the question of whether the retina itself holds mechanisms to reconfigure function-specific circuits in response to changing external influences.

Our work identifies a novel form of neural plasticity, present within the retina itself, which causes changes in retinal encoding for orientation tuning, dependent on the exposure of the animal to different visual environments during development. The modifications to the visual circuit occur across multiple levels, with changes in the retinal cell morphology and functionality, and innate animal behaviour. These findings reveal that the retina harbours mechanisms of plasticity that can significantly affect the interaction of the organism with its environment.

## Results

### Early exposure to different visual environments leads to orientation-specific altered amacrine cell morphology

To date, research on retinal plasticity has primarily focused on the effects of altering general sensory experience through deprivation but has overlooked the possible impact of changing salient visual features, such as orientation selectivity, which enables animals to recognise elongated or rectilinear shapes in visual space^17^. Retinal orientation selectivity in zebrafish is dependent on specific amacrine cells (ACs) with asymmetrically elongated dendritic arbours (i.e., type II and III in zebrafish)^29,34^. These retinal interneurons tune the output of orientation-selective (OS) retinal ganglion cells through orthogonally oriented feedforward inhibition, thus relating their structure directly with their function^29,30,35^. The development of visual responses in zebrafish is fast, starting in the first few days post fertilisation (dpf)^36^ and robust OS responses can be detected at 5 dpf^29,37^. To identify if early exposure of larvae to different visual environments affects the morphology and arrangement of ACs underlying orientation selectivity, animals were raised in narrow channels lined with differently arranged stripes of approximately 0.03 cycles per degree (CPD), a spatial frequency previously identified as the most effective for inducing OS responses^38^ between 0 and 5 dpf in zebrafish larvae. This environment provided a consistent and salient visual stimulus, presenting either horizontally or vertically oriented stripes to the fish, or no patterns (clear channels) as a control (Figure 1a).

**Figure 1:**
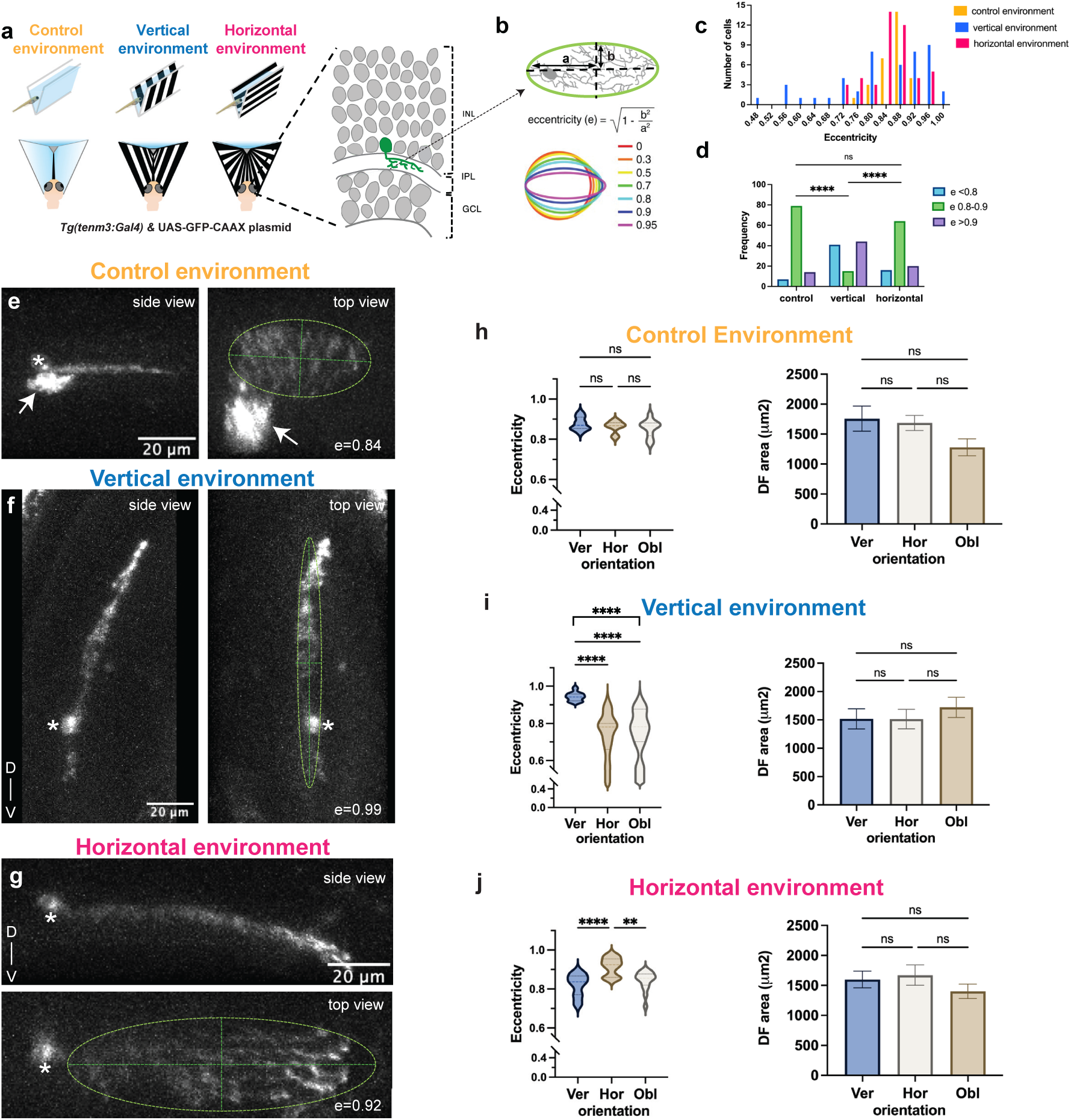
Exposure to different visual environments leads to changes in morphology of orientation-selective amacrine cells. **a.** Schematic of visually selective environment set up, with *Tg(tenm3:Gal4)* larvae, injected with a UAS-GFP-CAAX DNA plasmid raised in channels presenting either a control environment, vertical stripes, or horizontal stripes, from 72 hpf to 5 dpf and then assessed for sparse amacrine cell labelling. Top is a side-angle view, bottom is a point of view from the larvae’s perspective. Asymmetric type II amacrine cells were identified by dendritic arealisation and stratification in the inner plexiform layer (IPL). **b.** Eccentricity of the dendritic area was calculated by best fitting an ellipse on the cell and following the formula indicated. **c.** Distribution of eccentricities for larvae raised in control (yellow), vertical (blue) or horizontal (magenta) environments. **d.** Frequency of observed eccentricity in the three categories either falling below 0.8 (blue), between 0.8 and 0.9 (green) or >0.9 (purple). **e.** Side and top view of a type II amacrine cell from an animal raised in a non-patterned control environment. Asterisks indicates the soma, arrow indicates an additionally labelled type I cell. The green line indicates the ellipse used for eccentricity calculation. **f.** Side and top view of a type II cell from a larva raised in a vertical environment. Asterisks indicates the soma. **g.** Side and top view of a type II amacrine cell from an animal raised in a horizontal environment. Asterisks indicates the soma. **h, I, j.** Eccentricities (left graphs) or dendritic field areas (right graphs) of type II cells from animals raised in the three different environments, split into their placement along the retinal axes, either vertically along dorsal-ventral (Ver), horizontally along nasal-temporal (Hor) or oblique in any other direction (Obl). Graphs show mean and error bars ± s.e.m. n = 300 larvae for each condition. *(**P < 0.01, ****P < 0.0001;* One-way ANOVA followed by Tuckey multiple comparison test). D, dorsal; dpf, days post fertilisation; GCL, ganglion cell layer, hpf, hours post fertilisation; INL, inner nuclear layer, IPL, inner plexiform layer; V, ventral.

We then focused our analysis on OS type II ACs. Cells were fluorescently marked through sparse labelling using injection of *UAS:eGFP-CAAX* DNA plasmids into *Tg(tenm3:Gal4;mitfa-/-;roy+/-;alb+/-)* incross-derived embryos, which leads to the random labelling of several different AC subtypes, positive for the molecular marker teneurin-3 (tenm3), including type II ACs^29^ (Methods). Individual AC subtypes were identified based on their general shape and stratification in the inner plexiform layer^29^. The analysis focused on dendritic arborisation field (DF) area, the eccentricity (e) of the arborisation field and the directionality of the arborisation field (orientation) in relation to the eye axes for OS type II ACs (Figure 1b and Supplemental Figure 1b,c).

The eccentricity of type II ACs from fish raised in control channel conditions (n = 29) ranged between 0.76 and 0.92, with an average of e = 0.86 ± 0.03, which is consistent with values from larvae raised in normal Petri dishes, exhibiting an average of e = 0.86 ± 0.05 (n = 29, Figure 1c-e). However, when animals were raised in the horizonal or vertical environments, we observed in both cases a general broadening of the eccentricity distributions, ranging from 0.45 up to 0.99, (Figure 1c,d), while their dendritic field areas stayed the same (Supplemental Figure 1d). We then investigated if eccentricity is correlated with the alignment of the asymmetric dendritic fields of the type II cells along the retinal axes in the eye, either vertically (dorsal-ventral), horizontally (nasal-temporal) or obliquely (any other orientation) (Supplemental Figure 1b,c). In control conditions, we found no difference in eccentricity across the different dendritic field orientations (Figure 1h left, *P=0.4732*). However, when analysing cells from the vertically-raised animals, we surprisingly found that all vertically oriented cells had a significantly increased eccentricity (e = 0.94 ± 0.02, n = 19) compared to the horizontally (e = 0.74 ± 0.12, n = 16 (*P<0.0001*)) or obliquely positioned cells (e = 0.75 ± 0.12, n = 11, (*P<0.0001*)) (Fig 1f,i left). In turn, in retinae of horizontally-raised animals (n = 45), we found that the horizontally oriented cells showed a significantly increased eccentricity (e = 0.92 ± 0.05, n = 13) compared to the vertically (e = 0.82 ± 0.06, n = 16, *P<0.0001*) and obliquely oriented (e = 0.84 ± 0.05, n = 16, *P<0.0015*) cells (Fig1g,j left). Furthermore, when focusing on the cells showing lower eccentricity compared to the average value, we found that for animals raised in vertical stripes, all cells are oriented either horizontally or obliquely (Fig 1i, left), while for animals raised in horizontal stripes, the least eccentric cells are in the vertical and oblique category (Figure 1j left). In contrast, none of the vertically, horizontally, or oblique arranged cells exhibited a change in their dendritic field area (Figure 1h-j, right).

In summary, our morphological analysis discovered that raising zebrafish in vertically or horizontally patterned environments drastically changes the morphology of type II ACs. The cells with dendritic field extensions parallel to the presented pattern exhibit a significant elongation (higher e) while cells orthogonal or oblique to the presented pattern show a broadening (lower e). Since type II ACs are critical for generating orientation selectivity in the functional retinal output by OS retinal ganglion cells, we next investigated if developmental exposure to the differently striped environments would also alter the functional responses of retinal output when presented with different visual stimuli.

### Orientation-selective retinal output is dependent on developmental visual environment

To explore the functional plasticity of retinal circuits, we examined retinal ganglion cell (RGC) output responses in *Tg(isl2b:Gal4;UAS:SyGCaMP3;mitfa^-/-^)* zebrafish larvae at 5 dpf. In this line, all RGCs express the calcium indicator GCaMP3, fused to the presynaptic protein synaptophysin^33^. This allowed us to capture population visual responses through calcium imaging of RGC axon terminals in the optic tectum. Animals were raised in the different visual environments from 0-5 dpf. Awake larvae were then immobilized in small drops of agarose and presented with moving gratings in 12 different orientations to one eye, with visual responses recorded in RGC terminals present in the contralateral tectum (Figure 2a). Voxel-wise analysis allowed isolation of visually responsive voxels and determination of OS and direction-selective (DS) voxels. Subtypes of responses were identified via Gaussian distributions and grouped to preferred angles^29^ (Methods).

**Figure 2:**
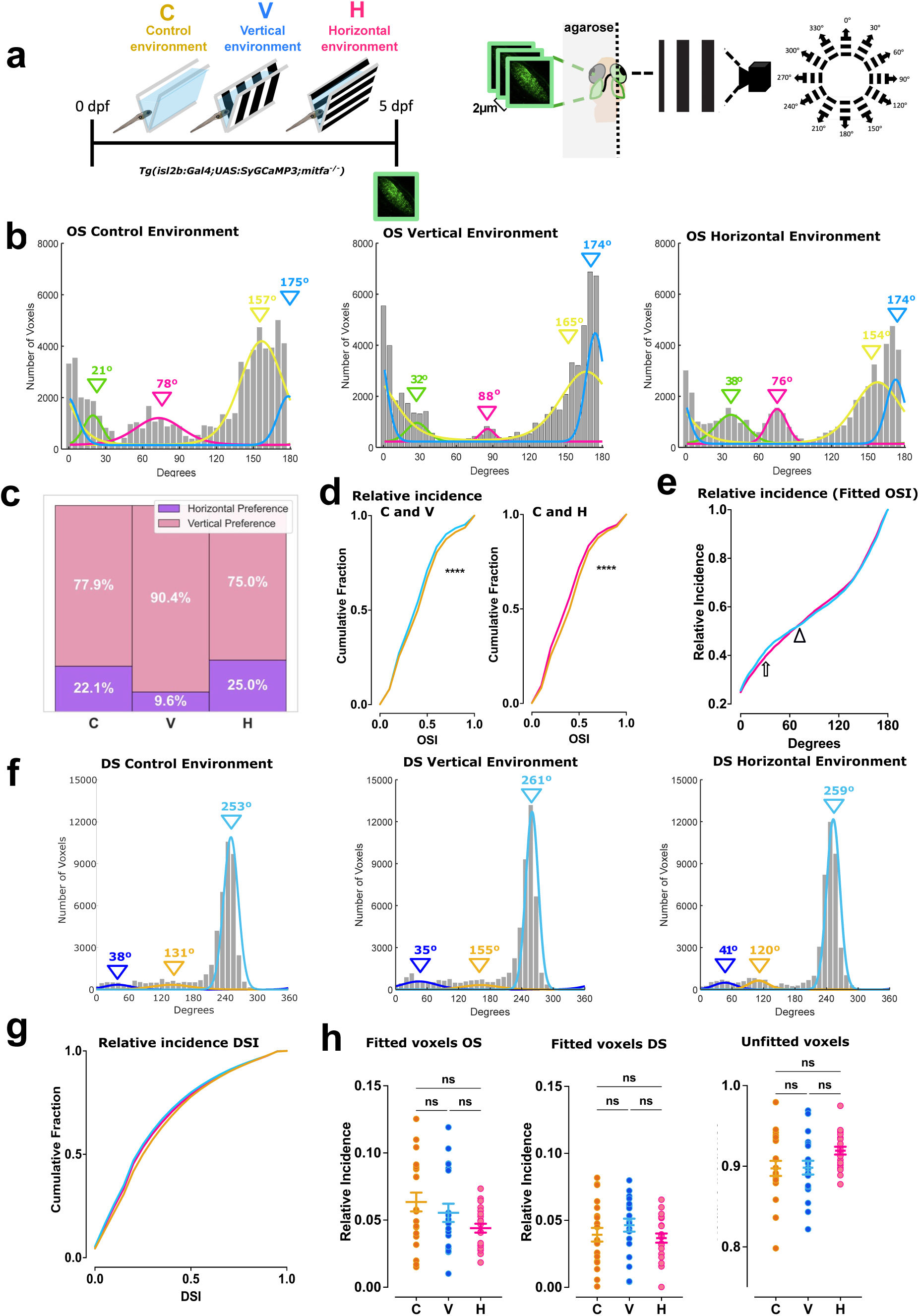
Exposure to different visual environments leads to changes in orientation-selective functional retinal output. **a.** Schematic of visually selective environment set up, with *Tg(isl2b:Gal4;UAS:SyGCaMP3)* larvae raised in channels presenting either a control environment (C), vertical stripes (V), or horizontal stripes (H) from 0 to 5 dpf and then assessed by calcium functional imaging of RGC terminals expressing SyGCaMP3 (green). Larvae are shown a stimulus of black and white stripes of various orientations moving in different directions, while recordings are taken from three Z-planes in the contralateral tectum. **b.** Histograms summarising the incidence of preferred angles for identified OS voxels in control larvae (n= 20), and those raised in a visually selective environment of vertical stripes and horizontal stripes (n = 20 each). Overlaid curves are the fitted Gaussian distributions for each OS subtype. **c.** Relative proportions for orientation selective retinal output, based on total voxels between 0° – 15° and 150° – 180° (vertical preference), and between 50° – 100° (horizonal preference) recorded in individual animals from control, vertical and horizontal environments. **d.** Cumulative distributions of OSI values (R^2^ > 0) across voxels with DSI < 0.5 for vertical (blue, left) and horizontal (magenta, right) versus controls (yellow) *(****P < 0.0001*, two-sample Kolmogorov-Smirnov test). **e.** Relative frequency of OSI responses across degrees for animals from vertical (blue) and horizontal (magenta) environments. Arrow indicates region with higher vertical incidence, arrowhead indicates crossing point at horizontal orientations. **f.** Histograms summarising the incidence of preferred angles for identified DS voxels in control larvae (n = 20), and those raised in a visually selective environment of vertical stripes and horizontal stripes (n = 20). Overlaid curves are the fitted Gaussian distributions for each DS subtype. **g.** Cumulative distributions of DSI values (R^2^ > 0) across voxels with OSI < 0.5 in vertical (blue) and horizontal (magenta) and controls (yellow). *\*\*\*\*P < 0.0001*, Kruskal-Wallis test. **h.** Relative frequency of OS, DS, and unfitted voxels per fish (n = 20 each condition). Criteria used to identify DS and OS voxels are reported in the methods. Graphs show mean and error bars ± s.e.m. over fish. (Brown-Forsythe ANOVA test with Dunnett’s T3 multiple comparisons test). dpf, days post fertilisation; C, control environment; DS, direction selective; H, horizontal environment; OS, orientation selective; OSI, orientation selectivity index; V, vertical environment.

Larvae raised in control channels showed the four innate peaks of OS subgroups within the RGC output corresponding to responses to vertical orientations (175°), horizontal orientations (78°), and two diagonal orientations (21° and 157°), consistent with previous findings^37,39^ (Figure 2b, left). However, for larvae raised in a vertically restricted environment, we observed a pronounced change in OS responses, with an increase in activity to stimuli of vertical orientations (175°), a decrease in response to horizontal stimuli (86°), and a slight shift in the two diagonal peaks towards the vertical orientations (32° and 165°) (Figure 2b, middle) In contrast, larvae raised in a visually selective environment of only horizontal stimuli, show a decrease in activity to stimuli of vertical orientations (174°) in comparison to vertical stimuli, an increase in response to horizontal stimuli (76°), and a shift in the two diagonal peaks towards the horizontal orientations (38° and 154°) (Figure 2b, right). We further find that the total voxel responses recorded within an orientation window either in the vertical range (0° -15° and 155° – 180°) or horizontal range (50° – 100°) is skewed towards vertical preference for animals raised in a vertical environment, or horizontal preference for horizontal environment, respectively (Figure 2c, Supplemental Figure 2). These findings strongly support the evidence of early environmental influence on functional retinal orientation selectivity tuning. Larvae raised in visually restricted environments exhibited a higher cumulative fraction of OS voxels compared to controls suggesting that their visual processing is more selective and finely tuned (Figure 2d). Analysing the relative incidence of the fitted OSI across patterned angles, we detect that the horizontally raised animals exhibit first a lower voxel selectivity (Figure 2e, arrow), but then meet the vertical incidence at the point of the horizonal peak responses (Figure 2e, arrowhead).

In contrast, we found no changes in DS responses for larvae raised in a visually selective environment of either horizontal or vertical stripes, with population size and relative proportions of DS subtypes continuous across all contrasting environments (Figure 2f,g). This suggests that DS subtypes are not affected by early visual experiences and their establishment remains constant during development, a finding consistent with previous research that observed no change in DS responses, even after dark-rearing of larvae^37^ or broad early silencing of neural activity^22^. Despite these shifts in OS responses, the overall proportion of identified OS or DS voxels remained stable relative to the total population of responsive voxels, with no significant difference between any of the conditions (Figure 2h). Additionally, the proportion of non-tuned RGC outputs did not change. This consistency suggests that exposure to different environments does not change the amounts of tuned and non-tuned retinal ganglion cells, but rather the encoding of specific OS information.

These findings show that retinal circuits are not merely passive filters of visual information but hold an adaptive capacity in response to the early visual environment during development. This plasticity is found for OS, but not for DS responses, therefore indicating the existence of feature-specific retinal plasticity.

### Altered retinal output persists after the removal of the visually selective environment

So far, we measured functional responses of RGCs in larvae at 5 dpf, directly after they were exposed to the different visual environments during their initial development. To investigate the persisting impact of this early occurring retinal plasticity, we focused on how orientation-selective circuits in the zebrafish retina continue to develop beyond early larval stages. Previous research has shown that by 7 dpf the projection and lamination of RGC axons from the retina to their tectal target have matured, with well-defined RGC subtypes^40^ and stable synaptic density along fully formed RGC branches^41,42^. To assess if the detected experience-dependent plasticity of functional RGC responses is still present at those mature stages, we placed the animals, after being exposed to differently orientated stripes between 0 and 5 dpf, into environments devoid of visual stimuli for an additional two days (Figure 3a). The walls of these environments were designed with a neutral grey, a midway contrast between the black and white stripes of the initial channels (Methods), to minimise additional visual input and isolate the impact of the early selective environments. Then, at 7 dpf, we assessed the visual responses using *in vivo* functional imaging as before. Remarkably, the shift in the proportion of OS responses observed at 5 dpf was maintained also at 7 dpf (Figure 3b, c). Specifically, larvae raised in vertical selective environments exhibited an increased proportion of vertically oriented RGC responses, accompanied by a decrease in horizontally oriented responses (Figure 3b middle, 3c). Conversely, larvae raised in horizontal environments showed an increase in horizontally oriented responses and a decrease in vertically oriented ones in comparison to visually selective vertical stimuli (Figure 3b right, 3c). For both conditions we again found significant changes of the relative incidence for OS voxels between animals raised in vertical and horizontal environments initially, with a subsequent joining of the values at the horizontal orientation (Figure 3d). However, as seen in larvae at 5 dpf, we did not find a decrease in the overall proportion of OS (Figure 3e, left) and DS (Figure 3e, middle) voxels, relative to the total population of responsive voxels, nor was there a change in the proportion of non-tuned RGC outputs (Figure 3e, right).

**Figure 3:**
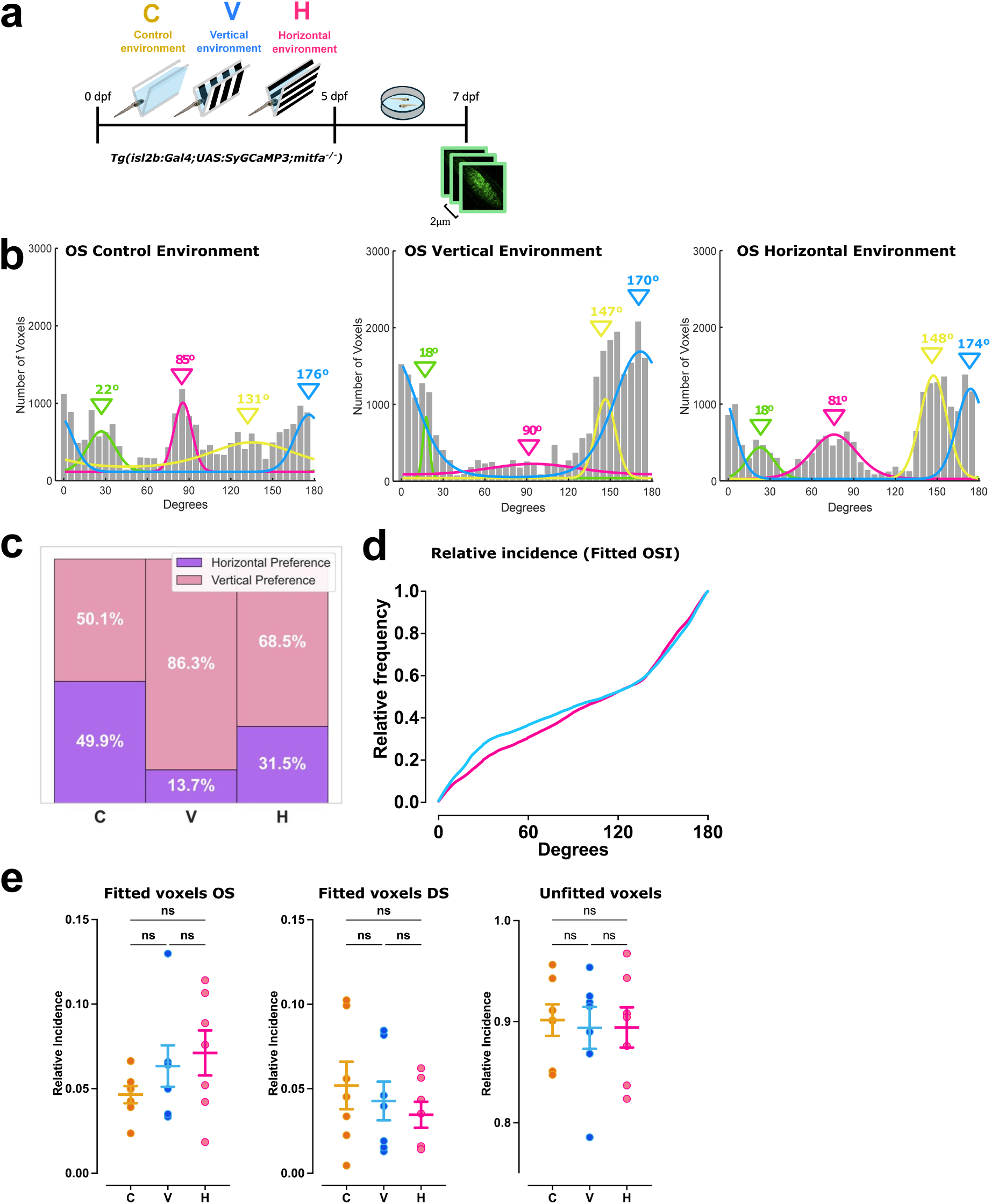
Functional changes in retinal orientation selectivity are long lasting. **a.** Schematic of experimental strategy, with *Tg(isl2b:Gal4;UAS:SyGCaMP3)* larvae raised in channels presenting either a control environment (C), vertical stripes (V), or horizontal stripes (H) from 0 to 5 dpf and then placed into a 35mm petri dish with a grey environment until 7dpf, before being assessed for retinal functional output. **b.** Histograms summarising the incidence of preferred angles for identified OS voxels in control larvae (n = 7), and those raised in a visually selective environment of vertical stripes and horizontal stripes (n = 7). Overlaid curves are the fitted Gaussian distributions for each OS subtype. **c.** Relative proportions for orientation selective retinal output, based on total voxels between 0° – 15° and 150° – 180° (vertical preference), and between 50° – 100° (horizonal preference) recorded in individual animals from control, vertical and horizontal environments. **d** Relative frequency of OSI responses across degrees for animals from vertical (blue) and horizontal (magenta) environments. **e.** Relative frequency of OS, DS and unfitted voxels per fish (n = 7 each condition). Criteria used to identify DS and OS voxels are reported in the methods. Graphs show mean and error bars ± s.e.m. over fish. (Brown-Forsythe ANOVA test with Dunnett’s T3 multiple comparisons test).

These findings highlight that early exposure to different visual environments shapes the functional output of the retina over a sustained period but does not alter the general ability to isolate different features from the visual scene. Here, we demonstrate that the plasticity of orientation-selective circuits in the zebrafish retina is not only robust but also enduring, with specific environmental influences shaping visual processing in a sustained manner beyond the initial stages of development.

We next aimed at investigating if the plasticity detected in the visual pathway influences the behaviour of the animals. For this, we established a novel behavioural paradigm, based on OS visual processing.

### Behavioural preference for differently orientated visual stimuli

The extraction of salient visual features within the retina is a core component of generating behaviours essential for survival. Detection of visual stimuli leads to feature-specific processing, which in turn triggers innate and learned responses^43^. Within zebrafish, this can include looming stimuli for escape responses^44^, prey capture^45^ and social interactions^46^. It is known that the visual environment can influence the animal’s behaviour^47–49^. However, current zebrafish paradigms usually quantify behaviour through moving gratings or dot-motion paradigms to mimic interactions with the natural environment^3,50,51^. Therefore, we implemented a new paradigm based on the presentation of non-moving grating patterns. This unique stimulus arrangement enables us to disentangle the relationship between the innate optomotor (OM) response and responses to a static pattern orientation as a salient visual feature.

We utilised a high-throughput virtual reality system, where stimuli are projected from below to freely swimming fish. The location and orientation of individual larvae are tracked in real-time, allowing us to keep stimuli fixed relative to the position and body orientation of animals (Figure 4a). This setup allowed us to present each eye of the free-swimming larvae with distinct and precisely controllable visual inputs to assess the impact of different stimuli to the animal’s behaviour.

**Figure 4:**
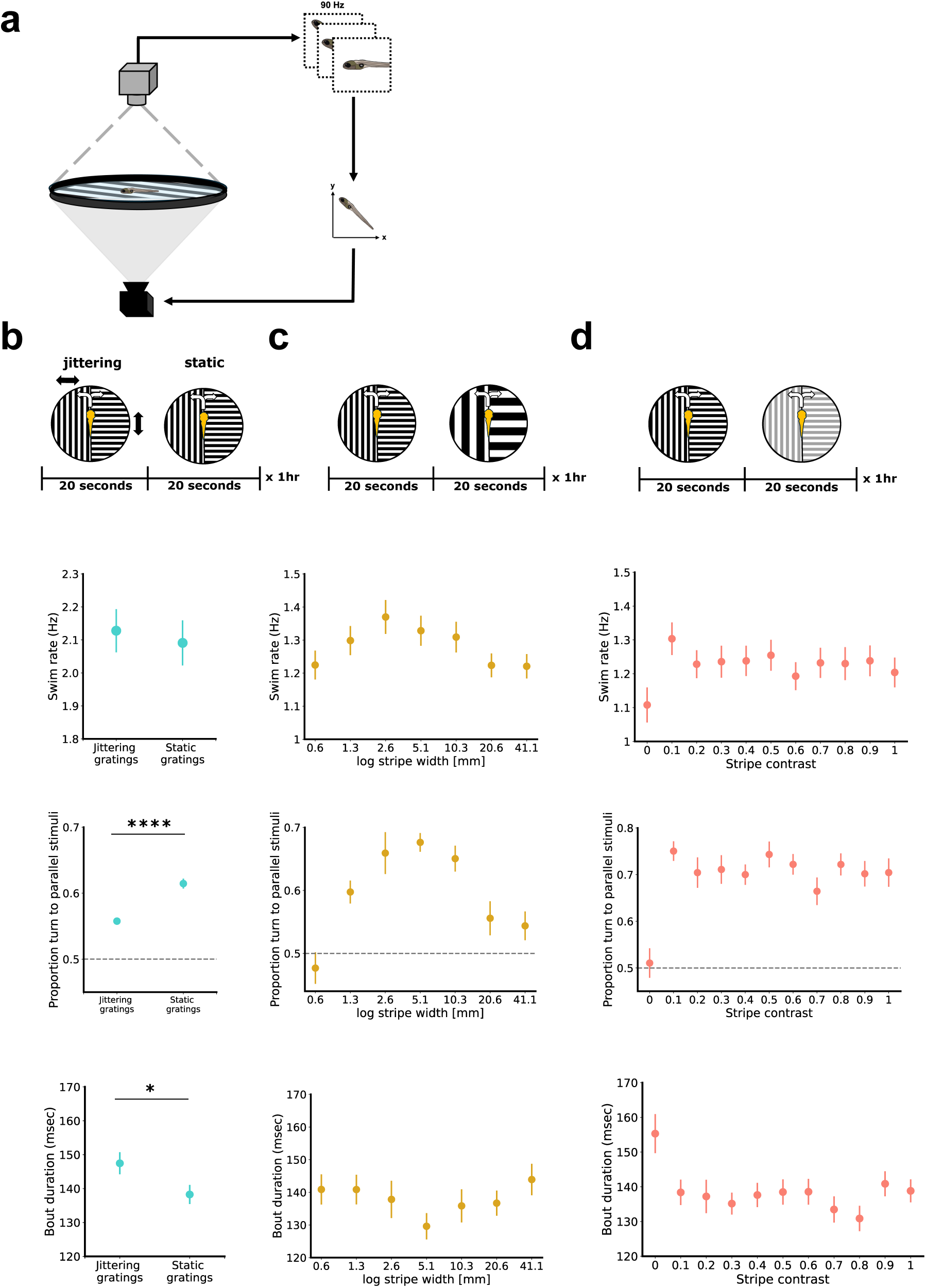
Novel visual behavioural paradigm uncovers innate animal preference to move towards parallel patterns. **a.** Schematic of closed-loop behaviour set up with stimuli presented from below to freely swimming larval zebrafish. Stimuli remain locked on to the larvae independent of position of the animal in the arena (12cm diameter). **b-d** Different stimulation parameters for behavioural assay: **b.** Jittering versus static: Animals experience a split-view configuration, with one eye given a stimulus parallel and the other eye a stimulus which is orthogonal to the orientation of the zebrafish. The stimulus changes between static gratings and jittering gratings randomly between each trial of 20 seconds (n=48 larvae). Significant differences between jittering and static stimuli were found for turn proportions (*\*\*\*\*P < 0.0001*) and bout duration (*\*P < 0.0351*). **c.** Stripe frequency: Animals experience a split-view configuration. The width of the gratings changes randomly between trials of 20 seconds, between 0.6mm to 41.1mm for the width of one black and one white grating (n = 32 larvae). Peak frequency response was found at 2.6mm, translated to a CPD of 0.03 when assuming the position of the fish. **d.** Stripe contrast: Stimuli presented to the freely swimming larvae from below in a split view configuration. The contrast between black and white lines changes as a function of overall contrast, from 0.0 (grey) to 1.0 (black and white) for one black and one white grating, indicated in example above graphs (n = 32 larvae). **b-d.** Frequency graphs indicate bout frequencies of larvae per second when shown different stimuli. Turning graphs show the proportion of turns towards parallel stimuli per all turns observed. Duration graphs indicate the measured bout duration of larvae when shown different types of stimuli. All error bars are mean ± s.e.m. over fish. P values are based on two-tailed t-test with Welch’s correction. Asterisks indicate significance *(*P < 0.05, ****P < 0.0001)*.

#### Behavioural responses to non-moving gratings

As we have identified that direction selectivity is not significantly influenced by the environment in which larvae were raised (Fig 2c), we wanted to isolate the effects of orientation selectivity from other visual processing mechanisms, such as OM responses based on direction selectivity. We therefore first compared the behavioural responses between standing gratings and gratings with small forward and backward motion (jittering) (Figure 4b). These jittering stimuli are designed to characterise potential innate behavioural outputs linked to OM output.

Firstly, we quantified a preference by measuring the proportion of total bout events per stimuli towards parallel stimuli. Our data revealed a clear preference for larvae to turn towards stimuli that are parallel to their body axes, over orthogonal stimuli (Figure 4b). This occurred for both jittering and static gratings, although larvae were significantly more likely to turn towards still stimuli aligned with their orientation, with the behaviour of turning towards parallel stimuli above chance (Figure 4b, mean = 0.5; moving gratings: mean = 0.5576, ± 0.0046; static gratings: mean = 0.6145, ± 0.0075, P < 0.0001). This indicates an innate response to orientation, in the absence of visual movement, and uncovers orientation selectivity to play a pivotal role in guiding zebrafish behaviour. Interestingly, we observed that this preference for parallel stimuli was robust, suggesting a potential innate bias towards parallel stimuli. Consequently, this may have implications on how zebrafish perceive and interact with their visual environment, with vertical based stimuli linked to depth perception and spatial understanding^52,53^.

We found no significant difference in overall behavioural outcomes, such as swim rate (Figure 4b), speed, interbout interval, or bout length (Supplemental Figure 3a) between jittering and still gratings. However, a slight increase in bout duration was noted with dynamic gratings (Figure 4b) (jittering gratings = mean: 147.5 msec, ± 3.2474; still gratings = mean: 138.2 msec, ± 2.8226), possibly due to a natural OM response through receiving visual feedback.

#### Behavioural responses to stripe width

We further examined how variations in stripe width influenced zebrafish behaviour, with previous behavioural paradigms using a range of widths from 1-20mm^28,54^. Here, we presented still gratings through a split-view paradigm with changing widths (Figure 4c). Animals displayed a strong preference for specific stripe widths, with a swim rate of bouts peaking at a stripe width of 2.6mm (Mean: 1.3699, ± 0.0512) for one black and white stripe (Figure 4c). Beyond this optimal stripe width, behavioural responses plateaued, particularly for larger widths of 20.6mm (mean: 1.2236, ± 0.0359) and 41.2mm (mean: 1.221, ± 0.0368), possibly due to the stripes becoming too large to be perceived by the larvae. Stripe width further influenced the animal’s preference for turning towards parallel stimuli (Figure 4c), following a slight right-skewed Gaussian distribution with a non-significant peak at 5.1mm (mean: 0.6763, ± 0.0149). This finding aligns with previous research on orientation tuning in zebrafish, which identified a similar optimal cycle per degree for eliciting the highest orientation-selective responses^38,55^, when assuming the location of the larvae within the arena.

Additionally, stripe width impacted bout duration (Figure 4c), with the shortest bout duration observed at a width of 5.1mm (mean: 129.6 msec, ± 4.0343). This suggests a range between 2.6mm and 5.1mm as the ideal stripe width for eliciting the most robust behavioural responses. Additional parameters, such as bout speed, interbout interval and bout length (Supplemental Figure 3b) confirmed an optimal stripe width between 2.6mm and 5.1mm. The impact of stripe width on these behavioural metrics underscores the importance of orientation processing in guiding zebrafish behaviour, highlighting the role of specific visual features in shaping innate responses.

#### Behavioural responses to stripe contrast

Lastly, we assessed the effects of stripe contrast on behavioural outputs (Figure 4d). Varying contrast had minimal impact on the behavioural properties of the larvae. All measured behavioural metrics, including frequency of bouts, proportion turn to parallel stimuli, and bout duration, all plateaued at a contrast level of 0.2 (Fig 4d). No significant change of behaviour was observed in any of these metrics as contrast increased, with frequency of bouts stabilizing at 1.2286 Hz (±0.0413), and the proportion turn to parallel at 0.7042 (±0.0325).

Similarly, additional behavioural outputs, such as speed and bout length (Supplemental Figure 3c), were not impacted with increasing contrast. Importantly, as with stripe width, the interbout interval remained consistent across different contrast levels (Supplemental Figure 3c), suggesting that the periods of rest between bouts were unaffected by changes in visual contrast, possibly suggesting no change in internal states through differing visual contrasts^56^.

In conclusion, our results demonstrate that non-moving gratings with an optimal stripe width and contrast effectively elicit robust OS responses in zebrafish larvae. These findings establish a new paradigm for investigating the impact of salient visual features on behaviour, independent of innate optomotor responses. This approach provided a foundation for further exploration of retinal plasticity and its role in shaping the neural mechanisms underlying visual perception and behaviour.

#### Visual experience induces behavioural plasticity

Environmental exposure is known to impact cognitive abilities in various species^57^. Previous research indicates that exposure to certain visual forms can enhance sensitivity within an OM response behaviour^28^. However, using our new paradigm, it was possible to directly assess how environmental structure influence behavioural output of zebrafish larvae, in the context of orientation selectivity. Although prior studies have hinted at such connections in humans, linking environmental experience or stimuli to higher-order visual processing and behaviour^9–11^, experimental constraints have resulted in limited understanding of how individual visual circuits can impact behavioural output.

Based on the insights from our novel visual stimulation paradigm, we designed a visual stimulus aimed at eliciting orientation-based behavioural responses while differentiating from innate OM responses (Figure 5a). The goal was to implement stimuli that could effectively drive locomotion (bouts) in zebrafish larvae, isolating the specific effects of orientation selectivity on innate behaviour.

**Figure 5:**
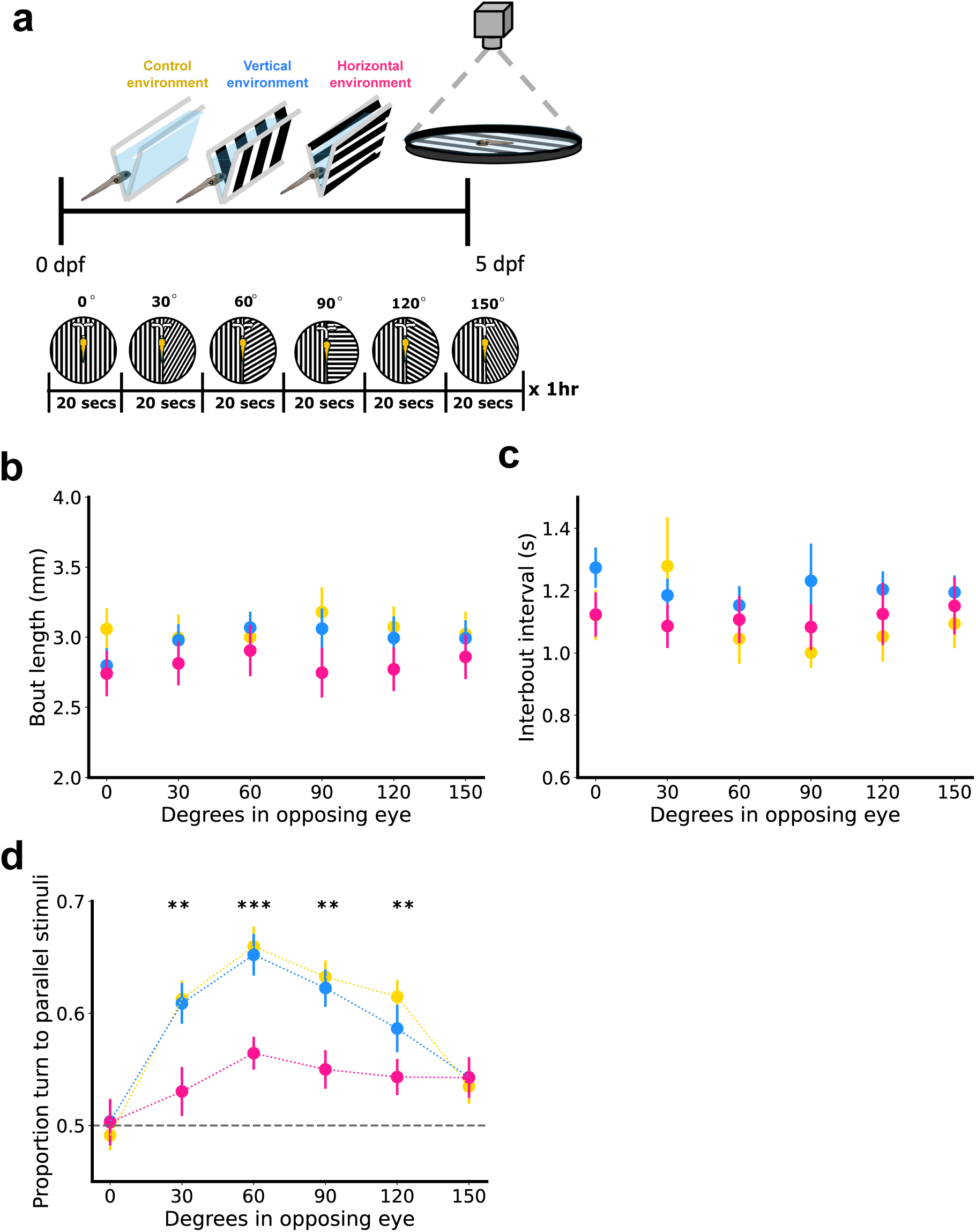
Early exposure to different visual environments leads to changes in animal behaviour. **a.** Schematic of experimental strategy, with larvae raised in channels presenting either a control (C) environment, vertical stripes (V), or horizontal stripes (H) from 0 to 5 dpf, then placed into the closed-loop behavioural arena. Stimuli remain locked on to the larvae independent of position of the animal in the arena (12cm diameter). One eye is given a stimulus always parallel to the orientation of the zebrafish, while the other eye receives a stimulus with changing orientations in random 30-degree steps between 0 and 150 degrees. **b.** Average bout length per stimulus when shown parallel stimuli in one eye and stimuli of changing angles in the opposing eye. **c.** Time between bouts (interbout interval) when shown parallel stimuli in one eye and stimuli of changing angles in the opposing eye. **d.** Proportion of turning towards parallel stimuli when shown parallel stimuli in one eye and changing stimuli in the opposing eye (30 degrees: \*\**P = 0.0016* (C vs H), \*\**P = 0.0042* (V vs H); 60 degrees: \*\*\**P = 0.0002* (C vs. H), \*\**P = 0.0011* (V vs. H); 90 degrees: \*\**P = 0.0015* (C vs. H), *\*\*P = 0093* (H vs. V); 120 degrees, \*\**P = 0.0073* (C vs H)). **b-d.** n = 64 for control environment raised larvae, n = 56 for vertical environment raised larvae, n = 72 for horizontal environment raised larvae. All error bars are mean ± s.e.m. over fish. (*\*\*P < 0.01, ***P < 0.001*, 2-way ANOVAs with multiple comparisons and corrected with Tukey test).

Zebrafish larvae were raised in visually selective environments from 0-5 dpf before their behaviour was tracked at 5 dpf using our split-view closed-loop system (Figure 5a). One eye consistently received a stimulus parallel to the larva’s orientation, while the other eye was exposed to stimuli varying in orientation in 30° steps between 0° (parallel to body axis) to 150°. We found no observable difference in the length of larval bouts, regardless of whether they were raised in horizontal, vertical, or control environments (Figure 5b). Similarly, the interbout interval remained consistent across all different early environmental experience conditions (Figure 5c), suggesting their basic swim characteristics were sustained.

We then asked whether the early exposure to different visual environments had a significant impact on the preferential response of the animal to the visual stimuli (Fig 5d). In control larvae, the preference to turn into the parallel stimuli resulted in a distinct tuning curve, with a peak-likelihood of turning towards parallel stimuli when the opposing eye was exposed to 60° oriented stimuli (mean = 0.6597, ± 0.0176). This pattern was mirrored in larvae raised in a vertically striped environment with a peak likelihood towards parallel stimuli at 60° in the opposite eye (mean = 0.6522, ± 0.0185). While there was a slight decrease in the likelihood of turning towards parallel stimuli at 120°, the overall shape of the distribution was preserved. However, notably, this OS behaviour was not maintained in larvae raised in a horizontal environment. Here the proportion of turning towards parallel stimuli rapidly decreased towards chance levels (mean = 0.5), with larvae significantly less likely to turn towards the parallel stimuli (opposite eye at 60°, mean = 0.5645, ± 0.0146). In addition, horizontal raised larvae showed no orientation tuning within the distribution between 30° and 150° (Figure 5d). This shift in increased likelihood towards chance strongly suggests that the environment in which the larvae were raised directly influences their behavioural output, particularly in response to orientation-specific visual stimuli. Assessment of bout speed, frequency and duration in this paradigm showed that larvae raised in a horizontal environment generally turn slower but more often, while the bout duration stays the same (Supplemental Figure 4a-c).

These findings provide compelling evidence for a novel form of plasticity, linking environmental conditions during early development to changes in behaviour and therefore highlight the importance of early visual patterns to shape the later visual functionality and the resulting behavioural output in zebrafish larvae.

## Discussion

Vision is vital for navigating through and interacting with our environment, and the capacity for neural circuits to adapt in response to sensory experiences is a fundamental requisite for appropriate behaviour^15,17–20,58^. In humans, this adaptation has been suggested to impact higher-order processes such as size perception^9^, object recognition^10^, and visuo-spatial processing^59^. However, sensory plasticity has traditionally been considered a phenomenon only expected outside the retina, with modifications to the retinal output only occurring to non-specific features in form of extensive visual deprivation^26,37,60–62^. Here, we challenge this long-held assumption, revealing that the sensory organ itself possesses a previously unrecognised form of plasticity for salient features, namely orientation selectivity, which is initially processed within the retina^29,30,34,63,64^. This remarkable capacity for feature-specific, experience-dependent plasticity is characterised by both morphological and functional changes within the retina when the animal is presented with different visual environments during development. We find that asymmetric amacrine cells underlying retinal orientation selectivity^29^ change their morphology in a visual experience-dependent manner. We further show that the functional OS retinal output is enhanced for the orientations presented to the animal early in development. By isolating retinal output, we minimised the influence of non-visual sensory contributions that can impact processing in the brain ^65,66^ and discovered that the retina itself can dynamically reshape its circuitry to adapt to environmental demands. This plasticity is specific to orientation-selective circuits, with direction-selective circuits remaining unaffected, underscoring the complexity between the different parallel visual channels^22,30,32,67,68^.

Importantly, so far it has been difficult to demonstrate behavioural consequences in animals after experience-dependent plasticity through the presentation of different visual stimuli. Using a novel visual behaviour paradigm specifically suited to assess the contribution of orientation selectivity, our results demonstrate that early developmental exposure to different visual environments can lead to observable changes in behaviour, thus linking sensory experience directly to a measurable behavioural output. This causal association accentuates the broader impact, uncovering a crucial role in shaping the integration of specific salient information and the organism’s interaction with its environment.

Our findings not only emphasise the retina’s role in selectively processing different types of visual information^40,69,70^ but also show its ability to actively adapt to the visual environment in form of cellular, functional, and behavioural changes. This plasticity likely serves a critical function, optimising the organism’s competency to process and respond to statistics of the local environment, which in turn could influence higher-order visual processing. These findings raise broader considerations, as the ability to elicit plasticity and habituation during development is crucial for learning and memory, with failure to do so associated with several visual and neurodevelopment disorders^7,71,72^. Our work sets the basis to approach further important challenges to determine the existence of critical windows for retinal plasticity and the impact on wider neural circuitry to fully understand the underlying mechanisms of how sensory input affects animal behaviour.

Overall, the implications of this retinal plasticity are profound, suggesting that the initial stages of sensory processing are more malleable than previously thought, with potential ramifications for understanding the neural basis of visual perception.

## Methods

### Zebrafish

Zebrafish husbandry and experiments were approved by the Animal Welfare and Ethics Review Body at KCL and conducted under the project license PP7266180 to R.H. Embryos were placed into filtered fish facility water (mains purified through reverse osmosis, reconstituted to pH7 with sodium bicarbonate and conductivity of 535μS with Tropic Marin Sea Salt) and placed into either 7cm Petri dishes or triangular prism channels (depending on experiment). There were no more than 50 larvae per Petri dish, or 3 larvae per channel. Zebrafish were maintained in an incubator at a temperature of 28.5°C on a 14-hour ON/10-hour OFF light cycle. For experiments on larvae over 5 days, larvae were fed live rotifers as in line with the KCL Fish Facility feeding protocol (1.5ml rotifer solution per 5 fish). Functional imaging experiments were performed on the transgenic line *Tg(isl2b:Gal4;UAS:SyGCaMP3;mitfa^-/-^),* while behaviour experiments were performed on the AB wildtype line (ZIRC, 5^th^ Generation). All experiments were performed either at 5 dpf or 7 dpf, at which age sex cannot be determined.

### Visual environment set up

The visually selective environments were created using a 12-channel reservoir made of PET-G with a pyramidal base (Thames Restek). Individual channels measure 7mm (w) x 10mm (h) x 70mm (l) with each holding 2ml of water. Up to 3 larvae were placed into each triangular based pyramidal channel from 0-5 dpf. Stripes were black and white opaque, either horizontal or vertical, and covered the two congruent sides of the triangular channel. For larvae raised in a visually deprived environment, the dish walls and base were an opaque grey with a printed pixel value of 128. This is at 50% contrast intensity between the black and white stripes, which were at printed pixel values of 0, and 225, respectively. Reservoirs were placed into a square bioassay dish (Corning) and lids were covered with frosted vinyl to remove non-specific visual stimuli from above. Idiosyncratic stripes were created using Photoshop (Adobe) at a width of 0.66mm, which correlated to a spatial frequency of 0.03 cycles per degree (c/°), identified in previous studies as the optimum width for orientation selective visual responses in zebrafish larvae^38^. Once removed from their visual environment, larvae were placed into a light-proof box to minimise exposure to visual stimuli before experimentation began.

### Functional imaging experiments in larval zebrafish

For *in vivo* imaging, non-anaesthetised larvae were screened at 5 dpf for strong SyGCaMP expression in the tectum. Positive larvae were immobilised in either 2% (5 dpf) or 3% (7 dpf) low melting point agarose prepared in Danieau solution. Larvae were mounted dorsal side up and on the edge of a raised glass platform, with their head pointing to the right. Agarose was then cut with a scalpel to free the right eye and provide an unobstructed view of the projection screen. Mounted larvae were then placed into a black box and left to acclimate for 30 minutes to reduce drift and movement of the larvae during recording sessions.

Larvae watched visual stimuli of moving gratings in the right eye while RGC projections were recorded in the contralateral tectum. Stimuli was created through a custom written LabView and MATLAB (MathWorks) code running through a ViSaGe stimulus presenter (Cambridge Research Systems) and presented laterally using a Mini C800S Smart DLP Projector (Toumei). Functional imaging was conducted on a LSM 710 confocal microscope with a spectral detection scan head and 20x/1.0 NA water-immersion objective (Zeiss). A time series of 1200 frames was obtained at a rate of 4.1Hz and a resolution of 0.415×0.415μm (256×256 pixels). Each larva was recorded at three planes in the optic tectum across 2μm intervals. Stimuli was comprised of moving black and white gratings of 12 different orientations at 30° intervals. The long axis of the bar was orthogonal to the direction of motion, and each stimulus was presented on the screen in a randomised order for 8 seconds, with a white blank interval of 10 seconds between each stimulus to allow GCaMP signals to return to baseline.

### Processing of functional imaging data

Preliminary analysis was based on previous scripts^33^. Time-series slices were realigned and corrected for motion through a rigid body algorithm (SPM12, UCL). Images were registered with the median kernel size of 1 voxel (0.415 μm) and spatially smoothed using the Gaussian smoothing kernel of 2 voxels (0.83 μm) to remove noise and improve the noise-to-signal ratio. Cubic spline interpolation of the inter-epoch interval was used to correct the low-frequency drifts in baseline. Anatomical reference images were created, and signal intensity changes of voxels were analysed over each epoch-interval for different orientations. Subtypes of direction selectivity (DS) and orientation selectivity (OS) voxels were identified through thresholding of OS = DSI < 0.5 and OSI > 0.5. DS = DSI > 0.5 and OSI < 0.5.

### Analysis of relative changes from functional imaging data

From all voxels that surpassed thresholding (OSI > 0.5), all fitted OSI angles were taken and converted from radians into degrees between 0° - 180°. Analysis of vertical and horizontal preference was done on both individual fish and the whole population. Vertical peaks for each population are taken as the degrees between 0° to 15° and 150° to 180°. Total OS voxels are selected in 5 degree bins for the initial histogram. The histogram of the data was shifted to incorporate these two vertical voxel populations within one peak. Then, a best fit curve was generated matching the values of the histogram. Peaks were calculated for the shifted histogram and a designated vertical and horizontal peak was determined (peak within 0° to 15° and 150° to 180° for the vertical peak, and 50° to 100° range for the horizontal peak). The range was established based on histograms of individual fish (N = 60) and identifying vertical and horizontal peaks. The values of the peaks were used to calculate a vertical preference index for either a group or an individual fish using ∑(vertical voxels) / (∑(vertical voxels) + ∑(horizontal voxels)). A vertical preference index is calculated per fish and these values are transformed so the zero baseline represents cases where the vertical and horizontal voxel counts are equal. Larvae with an index value above zero show more vertical preference while those below zero show a more horizontal preference.

### Sparse stochastic labelling of Tenm3^+^ Amacrine cells

To randomly label *Tenm3*+ ACs, 1nL of UAS:eGFP-CAAX, DNA construct was injected into 1-2 cell-stage Tg(*tenm3:Gal4;mitfa^-/-^;roy^+/-^;alb^+/-^*) incross-derived embryos. Plasmid DNA was prepared using midiprep kits (Qiagen) and injected at a concentration of 30 ng/μl in ddH2O. This concentration was chosen to stochastically label up to 10 tenm3+ ACs, minimising the chances of AC overlap. Larvae were raised in 200μM PTU (Sigma) in Danieau solution to avoid pigment formation within the retina. UAS:eGFP-CAAX injected Tg(*tenm3:Gal4)* embryos were initially raised in 90 mm Petri dishes containing Danieau solution until 2 dpf. Then, embryos were screened at 3 dpf under fluorescence microscope for the presence of GFP signal in the retina. Only GFP+ larvae were selected and raised from 3 dpf to 5 dpf in different visual environments as outlined above.

### Confocal structural imaging

Confocal structural imaging of tenm3+ ACs was performed using a Zeiss LSM880 Axio Examiner equipped with a W Plan-Apochromat 20X/1.0 NA water-immersion objective. Larvae at 5 dpf were mounted in 0.3% agarose in 55 mm Petri dishes and anesthetised with 0.02% tricaine in Danieau solution. All larvae were mounted “Anterior to the left (AtL)”. Z-stacks were conducted of the whole eye for each screened larva. On average, 32-40 larvae were analysed during each imaging session and a total amount of 300 larvae was analysed per each environmental condition. Optical sections were typically obtained at 1 μm intervals through the Z axis and 1 AU pinhole aperture. Excitation light was provided by 488 nm multi-line laser and images were acquired at 0.49 × 0.49 μm resolution (512 × 512 pixels). Maximum intensity projections and 3D rotated images were generated using ImageJ.

### Morphological analysis

To define the cell type of each tenm3+ AC observed, the position of the cell body in relation to the arborisation field and the stratification of the arborization field within the IPL were used as key landmarks. Tenm3+ type II ACs were identified based on previous description^29^, and the dendritic field elongation, orientation, and placement was recorded. To quantify the dendritic field elongation of individual tenm3+ type II ACs, the eccentricity of dendritic arbour profiles [e = √(1 – b2/a2); a = length of the ellipse semi-major axis, b = length of the ellipse semi-minor axis] was calculated following ellipse fitting in ImageJ.

3D morphological reconstructions of individual tenm3+ type II ACs were obtained using the 3D project function in Fiji. Both arborization field’s eccentricity and area (in μm^2^) were measured using ImageJ in 3D rotated images. The relative frequency of each tenm3+ ACs type was finally calculated for each environmental condition by dividing the number of cells of each specific type (e.g. number of type II ACs) with the total amount of tenm3+ ACs (all 7 types) observed.

### 3D registration of type II ACs

A pipeline on MATLAB was developed to register and quantify the cell orientation of type II ACs from multiple animals to the same space. Registration was based on the position and orientation of the lens, and the relative position of the type II ACs to the lens. To segment the lens, the image was first up-sampled in the Z-axis, smoothened with a 3D Gaussian filter, brightened using contrast adjustment, and then binarized using thresholding. The long axis of the lens, defined as the line joining the centroids of the region of interest in each X-Y plane, was aligned to the Z-axis by translation and rotation of the image. Since the portion of the lens imaged varied across different animals, the centroid of the lens is estimated by fitting a circle to the back of the lens. The image is then translated such that the centre of the fitted circle is at the origin of the coordinate system. From the above translations and rotations steps, we obtain an affine transformation matrix. Due to the complex geometry and the possibility of overlapping cells, it was challenging to segment cells automatically. For this reason, the 3D coordinates of the cell body and both ends of the dendritic fields were obtained manually using Fiji/ImageJ. The corresponding affine transformation matrix was applied to map these coordinates into the same coordinate system. To adjust for the different sizes of eyes, the distance between the dendritic fields and the estimated centroid of the lens was then normalized to 1.

### Behaviour experiments in freely swimming larval zebrafish

Closed-loop behaviour paradigms in freely swimming larvae were conducted on an adapted system^50^. Larvae at 5 dpf were placed into custom-made circular acrylic dishes, measuring 120mm in diameter, 5mm in height, with a black rim and transparent base covered with diffusion paper. Experiments were conducted at 5 dpf, with larvae left in the dark behaviour room for 30 minutes before trials began to allow acclimatisation to the room. Dishes were filled with 50ml of filtered fish water, with one larva per dish. One computer was connected to 8 cameras and 4 projectors covering 8 dishes, allowing tracking of up to 8 larvae simultaneously. Stimuli were shown to the larvae using AAXA Neo-Pico projectors and the dishes were illuminated for the camera using infrared (940nm) light-emitting diode (LED) strip panels from below. Larvae were tracked using a high-speed CMOS camera (Basler acA2040-90um-NIR) with a zoom lens (Zoom 7000, 18-108mm) and an infrared filter. Live tracking was performed in real-time at a rate of 90fps using custom written behavioural tracking software on Python 3.11 and OpenCV4.1. Larval position and orientation were based on the overall shape of the fish, which were detected by the difference between each current frame and the average over the frames corresponding to the last 800 seconds and updated every 50 frames through background subtraction. Bouts were considered turns if they had an absolute orientation change larger than 2 °. Combinations of split-view stimuli were presented randomly to both eyes.

### Freely swimming paradigms

#### Still and moving gratings

Split-view stimuli with one eye shown parallel stimuli of 0°/180°, and the other shown horizontal stimuli of 90°/270°, with a randomly allocated offset pattern. Stripe size for each stimulus were a value of 1.92 mm for one black and white bar, and contrast was at 0.70. Gratings would either be static or move at a speed of 0.15 cm/s. Movement of gratings were so that the long axes were perpendicular to their direction of motion Each stimulus lasted for 20 seconds, with 90 trials per stimulus (1 hour duration) per fish, and water was kept at a temperature of approximately 26° continuously.

#### Stripe width

Split-view stimuli with one eye shown parallel stimuli, and the other shown horizontal stimuli, with a randomly allocated offset pattern. Stripe frequency for each stimulus ranged from 0.005 – 0.32 AU (0.005, 0.01, 0.02, 0.04, 0.08, 0.16, 0.32), equating to widths between 0.64 mm – 41.14 mm for one black and white bar, and contrast was at 0.7. This resulted in 14 stimuli combinations. Each stimulus had a contrast of 1 and lasted 20 seconds, with 13 trials per stimulus (1 hour duration) per fish. Water was kept at a temperature of approximately 26° continuously.

#### Contrast

Split-view stimuli with one eye shown vertical stimuli of 0°/180°, and the other shown horizontal stimuli of 90°/270°, with a randomly allocated offset pattern. Stripe size for each stimulus were a value of 1.92 mm for one black and white bar, and contrasts ranged from 0.0 – 1.0 with intervals of 0.1. Therefore, there are 20 stimuli combinations. Each stimulus lasted for 20 seconds, with 9 trials per stimulus (1 hour duration) per fish. Water was kept at a temperature of approximately 26° continuously.

#### Plastic responses

Larvae were raised in channels showing horizontal stimuli, vertical stimuli, or controls until 5 dpf, and then placed into the following closed-loop conditions: Split-view stimuli with one eye shown vertical stimuli of 0°/180°, and the other shown an angle between 0-180° in 30° intervals (30°, 60°, 90°, 120°, 150°), with a randomly allocated offset pattern. This resulted in 12 stimuli combinations. Stripe size for each stimulus was 2.6mm for one black and white bar, and with a contrast of 0.7. Each stimulus lasted 20 seconds, with 15 trials per stimulus (1 hour duration) per fish. Water was kept at a temperature of approximately 26° continuously.

#### Processing of behavioural data

Behavioural quantification is performed with tracking methods as described^50^. Swim events (bouts) were identified by periods of high activity in the rolling variance (50ms window size) of the body orientation. All behavioural experiments are pre-processed, with swim events dropped to exclude potential tracking errors. Events are dropped if their contour area is more than 2000 px, the interbout interval is more than 10 s, the average speed is more than 6 cm/s, the absolute orientation change more than 150 degrees, or the position within 0.5 cm from the wall. Such extreme events are considered tracking errors. In addition, a complete trial is dropped if these cases form more than 5% of all detected events in that trial. Each stimuli lasted 20 seconds, and the first 10 seconds of data was excluded to allow for the fish to adjust to the changing stimuli.

#### Statistics and reproducibility

All statistical analyses and tests were completed on Prism10 (GraphPad), MATLAB (R2014b/2023a, MathWorks), or Python. Descriptive statistics including normality were completed to choose the appropriate test. Statistical significance was set at p < 0.05 and sample sizes were chosen to provide a high statistical power of > 0.8.

For behavioural studies establishing visual stimuli and innate responses, we ran at least n = 32, in line with previous papers^50,73^. For studies quantifying the effect of plasticity on behaviour, we ran at least an n = 56 fish. These fish were chosen randomly within a Petri dish and came from between 2 - 6 parent batches per experiment, with batches collected and run on separate days. The processing script removes any trials where the animal does not move in the middle of the dish or could not be accurately tracked.

For functional imaging, we ran n = 20 (5 dpf), and n = 7 (7 dpf) per condition, recording from three z-planes imaged per larvae, as described before^29,37^. Data for each condition was collected from at least 2 (7 dpf) or 4 (5 dpf) different parent batches from multiple pairings. For structural imaging, an n = 300 larvae for each visual environment were used to identify tenm+ ACs .

#### Data availability

Raw data and computer code used to analyse data can be requested from the corresponding author.

## Acknowledgements

This work was supported by the MRC (MRC CNDD MR/W006251/1 to RH; MRC DTP MR/N013700/1 to PR), The Leverhulme Trust (RPG-2021-385 to RH), The BBSRC (BBSRC Int. Partnering Award BB/X512047/1 to RH, PR, AB, MC), Emmy Noether Program (BA 5923/1-1 to AB, MC), Zukunftskolleg Konstanz (AB, MC), and Boehringer Ingelheim Fonds (KS), and pilot work was funded by a University of Oxford Laidlaw Scholarship (PR). We thank Oscar Marin and Sam Cooke for their useful comments on the manuscript, and members of the team for their insightful discussions.

## Author Contributions

RH conceived the overall study and with help of PR and AB (behaviour), designed the experiments. PR (functional imaging & behaviour) and DM (cell morphology) performed all the experiments and analysed the data, with input from AB, KS, MC, YL, ZC. AB and MC helped to build the closed-loop system at King’s College London. AB supervised PR for initial training (behaviour) in Konstanz. PR provided the first draft and together with RH wrote the manuscript with input from all the other authors.

## References

1 Ahrens, M. B. et al. Brain-wide neuronal dynamics during motor adaptation in zebrafish. Nature 485, 471–477, doi:10.1038/nature11057 (2012).

2 Contier, O., Baker, C. I. & Hebart, M. N. Distributed representations of behaviour-derived object dimensions in the human visual system. Nat Hum Behav, doi:10.1038/s41562-024-01980-y (2024).

3 Kubo, F. et al. Functional architecture of an optic flow-responsive area that drives horizontal eye movements in zebrafish. Neuron 81, 1344–1359, doi:10.1016/j.neuron.2014.02.043 (2014).

4 Rao, S. et al. A direct and melanopsin-dependent fetal light response regulates mouse eye development. Nature 494, 243–246, doi:10.1038/nature11823 (2013).

5 Axelrod, C. J., Gordon, S. P. & Carlson, B. A. Integrating neuroplasticity and evolution. Curr Biol 33, R288–R293, doi:10.1016/j.cub.2023.03.002 (2023).

6 Jacobson, G. A., Rupprecht, P. & Friedrich, R. W. Experience-Dependent Plasticity of Odor Representations in the Telencephalon of Zebrafish. Curr Biol 28, 1–14 e13, doi:10.1016/j.cub.2017.11.007 (2018).

7 Ribic, A. Stability in the Face of Change: Lifelong Experience-Dependent Plasticity in the Sensory Cortex. Front Cell Neurosci 14, 76, doi:10.3389/fncel.2020.00076 (2020).

8 Ulanovsky, N., Las, L. & Nelken, I. Processing of low-probability sounds by cortical neurons. Nat Neurosci 6, 391–398, doi:10.1038/nn1032 (2003).

9 Doherty, M. J., Tsuji, H. & Phillips, W. A. The context sensitivity of visual size perception varies across cultures. Perception 37, 1426–1433, doi:10.1068/p5946 (2008).

10 Ueda, Y. et al. Cultural Differences in Visual Search for Geometric Figures. Cogn Sci 42, 286–310, doi:10.1111/cogs.12490 (2018).

11 Rivers, W. H. The measurement of visual illusion. Report of the British Association for the Advancement of Science, 818 (1901).

12 Barlow, H. B. Visual experience and cortical development. Nature 258, 199–204, doi:10.1038/258199a0 (1975).

13 Grubb, M. S. & Thompson, I. D. The influence of early experience on the development of sensory systems. Curr Opin Neurobiol 14, 503–512 (2004).

14 White, L. E. & Fitzpatrick, D. Vision and cortical map development. Neuron 56, 327–338, doi:10.1016/j.neuron.2007.10.011 (2007).

15 Blakemore, C. & Van Sluyters, R. C. Innate and environmental factors in the development of the kitten’s visual cortex. J Physiol 248, 663–716, doi:10.1113/jphysiol.1975.sp010995 (1975).

16 Crair, M. C., Gillespie, D. C. & Stryker, M. P. The role of visual experience in the development of columns in cat visual cortex. Science 279, 566–570, doi:10.1126/science.279.5350.566 (1998).

17 Hubel, D. H. & Wiesel, T. N. Receptive fields, binocular interaction and functional architecture in the cat’s visual cortex. J Physiol 160, 106–154 (1962).

18 Hubel, D. H. & Wiesel, T. N. The period of susceptibility to the physiological effects of unilateral eye closure in kittens. J Physiol 206, 419–436, doi:10.1113/jphysiol.1970.sp009022 (1970).

19 Hubel, D. H., Wiesel, T. N. & Stryker, M. P. Orientation columns in macaque monkey visual cortex demonstrated by the 2-deoxyglucose autoradiographic technique. Nature 269, 328–330, doi:10.1038/269328a0 (1977).

20 Kreile, A. K., Bonhoeffer, T. & Hubener, M. Altered visual experience induces instructive changes of orientation preference in mouse visual cortex. J Neurosci 31, 13911–13920, doi:10.1523/JNEUROSCI.2143-11.2011 (2011).

21 Strettoi, E., Di Marco, B., Orsini, N. & Napoli, D. Retinal Plasticity. Int J Mol Sci 23, doi:10.3390/ijms23031138 (2022).

22 Barabasi, D. L., Schuhknecht, G. F. P. & Engert, F. Functional neuronal circuits emerge in the absence of developmental activity. Nat Commun 15, 364, doi:10.1038/s41467-023-44681-2 (2024).

23 Sherman, S., Arnold-Ammer, I., Schneider, M. W., Kawakami, K. & Baier, H. Retina-derived signals control pace of neurogenesis in visual brain areas but not circuit assembly. Nat Commun 14, 6020, doi:10.1038/s41467-023-40749-1 (2023).

24 Feller, M. B., Wellis, D. P., Stellwagen, D., Werblin, F. S. & Shatz, C. J. Requirement for cholinergic synaptic transmission in the propagation of spontaneous retinal waves. Science 272, 1182–1187 (1996).

25 Ge, X. et al. Retinal waves prime visual motion detection by simulating future optic flow. Science 373, doi:10.1126/science.abd0830 (2021).

26 Cline, H. T. Activity-dependent plasticity in the visual systems of frogs and fish. Trends Neurosci 14, 104–111, doi:10.1016/0166-2236(91)90071-2 (1991).

27 Emran, F., Rihel, J., Adolph, A. R. & Dowling, J. E. Zebrafish larvae lose vision at night. Proc Natl Acad Sci U S A 107, 6034–6039, doi:10.1073/pnas.0914718107 (2010).

28 Xie, J., Jusuf, P. R., Bui, B. V. & Goodbourn, P. T. Experience-dependent development of visual sensitivity in larval zebrafish. Sci Rep 9, 18931, doi:10.1038/s41598-019-54958-6 (2019).

29 Antinucci, P., Suleyman, O., Monfries, C. & Hindges, R. Neural Mechanisms Generating Orientation Selectivity in the Retina. Curr Biol 26, 1802–1815, doi:10.1016/j.cub.2016.05.035 (2016).

30 Baden, T. et al. The functional diversity of retinal ganglion cells in the mouse. Nature 529, 345–350, doi:10.1038/nature16468 (2016).

31 D’Souza, S. & Lang, R. A. Retinal ganglion cell interactions shape the developing mammalian visual system. Development 147, doi:10.1242/dev.196535 (2020).

32 Kolsch, Y. et al. Molecular classification of zebrafish retinal ganglion cells links genes to cell types to behavior. Neuron 109, 645–662 e649, doi:10.1016/j.neuron.2020.12.003 (2021).

33 Nikolaou, N. et al. Parametric functional maps of visual inputs to the tectum. Neuron 76, 317–324, doi:S0896-6273(12)00806-9 [pii] 10.1016/j.neuron.2012.08.040 (2012).

34 Antinucci, P. & Hindges, R. Orientation-Selective Retinal Circuits in Vertebrates. Front Neural Circuits 12, 11, doi:10.3389/fncir.2018.00011 (2018).

35 Bloomfield, S. A. Orientation-sensitive amacrine and ganglion cells in the rabbit retina. J Neurophysiol 71, 1672–1691 (1994).

36 Easter, S. S., Jr. & Nicola, G. N. The development of vision in the zebrafish (Danio rerio). Dev Biol 180, 646–663, doi:10.1006/dbio.1996.0335 (1996).

37 Lowe, A. S., Nikolaou, N., Hunter, P. R., Thompson, I. D. & Meyer, M. P. A systems-based dissection of retinal inputs to the zebrafish tectum reveals different rules for different functional classes during development. J Neurosci 33, 13946–13956, doi:33/35/13946 [pii] 10.1523/JNEUROSCI.1866-13.2013 (2013).

38 Johnston, J. et al. A Retinal Circuit Generating a Dynamic Predictive Code for Oriented Features. Neuron 102, 1211–1222 e1213, doi:10.1016/j.neuron.2019.04.002 (2019).

39 Antinucci, P., Nikolaou, N., Meyer, M. P. & Hindges, R. Teneurin-3 specifies morphological and functional connectivity of retinal ganglion cells in the vertebrate visual system. Cell Rep 5, 582–592, doi:10.1016/j.celrep.2013.09.045 (2013).

40 Robles, E., Laurell, E. & Baier, H. The retinal projectome reveals brain-area-specific visual representations generated by ganglion cell diversity. Curr Biol 24, 2085–2096, doi:10.1016/j.cub.2014.07.080 (2014).

41 Meyer, M. P. & Smith, S. J. Evidence from in vivo imaging that synaptogenesis guides the growth and branching of axonal arbors by two distinct mechanisms. J Neurosci 26, 3604–3614, doi:10.1523/JNEUROSCI.0223-06.2006 (2006).

42 Niell, C. M., Meyer, M. P. & Smith, S. J. In vivo imaging of synapse formation on a growing dendritic arbor. Nat Neurosci 7, 254–260, doi:10.1038/nn1191 (2004).

43 Chen, P. & Hong, W. Neural Circuit Mechanisms of Social Behavior. Neuron 98, 16–30, doi:10.1016/j.neuron.2018.02.026 (2018).

44 Dunn, T. W. et al. Neural Circuits Underlying Visually Evoked Escapes in Larval Zebrafish. Neuron 89, 613–628, doi:10.1016/j.neuron.2015.12.021 (2016).

45 Bianco, I. H., Kampff, A. R. & Engert, F. Prey capture behavior evoked by simple visual stimuli in larval zebrafish. Front Syst Neurosci 5, 101, doi:10.3389/fnsys.2011.00101 (2011).

46 Kappel, J. M. et al. Visual recognition of social signals by a tectothalamic neural circuit. Nature 608, 146–152, doi:10.1038/s41586-022-04925-5 (2022).

47 Handegard, N. O. et al. The dynamics of coordinated group hunting and collective information transfer among schooling prey. Curr Biol 22, 1213–1217, doi:10.1016/j.cub.2012.04.050 (2012).

48 Sainsbury, T. T. J., Diana, G. & Meyer, M. P. Topographically Localized Modulation of Tectal Cell Spatial Tuning by Complex Natural Scenes. eNeuro 10, doi:10.1523/ENEURO.0223-22.2022 (2023).

49 Wei, D., Talwar, V. & Lin, D. Neural circuits of social behaviors: Innate yet flexible. Neuron 109, 1600–1620, doi:10.1016/j.neuron.2021.02.012 (2021).

50 Bahl, A. & Engert, F. Neural circuits for evidence accumulation and decision making in larval zebrafish. Nat Neurosci 23, 94–102, doi:10.1038/s41593-019-0534-9 (2020).

51 Naumann, E. A. et al. From Whole-Brain Data to Functional Circuit Models: The Zebrafish Optomotor Response. Cell 167, 947–960 e920, doi:10.1016/j.cell.2016.10.019 (2016).

52 Matthews, N., Meng, X., Xu, P. & Qian, N. A physiological theory of depth perception from vertical disparity. Vision Res 43, 85–99, doi:10.1016/s0042-6989(02)00401-7 (2003).

53 Yacoub, E., Harel, N. & Ugurbil, K. High-field fMRI unveils orientation columns in humans. Proc Natl Acad Sci U S A 105, 10607–10612, doi:10.1073/pnas.0804110105 (2008).

54 Kist, A. M. & Portugues, R. Optomotor Swimming in Larval Zebrafish Is Driven by Global Whole-Field Visual Motion and Local Light-Dark Transitions. Cell Rep 29, 659–670 e653, doi:10.1016/j.celrep.2019.09.024 (2019).

55 Preuss, S. J., Trivedi, C. A., vom Berg-Maurer, C. M., Ryu, S. & Bollmann, J. H. Classification of object size in retinotectal microcircuits. Curr Biol 24, 2376–2385, doi:10.1016/j.cub.2014.09.012 (2014).

56 Johnson, R. E. et al. Probabilistic Models of Larval Zebrafish Behavior Reveal Structure on Many Scales. Curr Biol 30, 70–82 e74, doi:10.1016/j.cub.2019.11.026 (2020).

57 Montalbano, G., Bertolucci, C. & Lucon-Xiccato, T. Cognitive Phenotypic Plasticity: Environmental Enrichment Affects Learning but Not Executive Functions in a Teleost Fish, Poecilia reticulata. Biology (Basel) 11, doi:10.3390/biology11010064 (2022).

58 Bauer, J. et al. Sensory experience steers representational drift in mouse visual cortex *bioRxiv* p. 2023.09.22.558966 (2023).

59 Gonthier, C. Cross-cultural differences in visuo-spatial processing and the culture-fairness of visuo-spatial intelligence tests: an integrative review and a model for matrices tasks. Cogn Res Princ Implic 7, 11, doi:10.1186/s41235-021-00350-w (2022).

60 Holtmaat, A. & Svoboda, K. Experience-dependent structural synaptic plasticity in the mammalian brain. Nat Rev Neurosci 10, 647–658, doi:10.1038/nrn2699 (2009).

61 Levelt, C. N. & Hubener, M. Critical-period plasticity in the visual cortex. Annu Rev Neurosci 35, 309–330, doi:10.1146/annurev-neuro-061010-113813 (2012).

62 Wiesel, T. N. & Hubel, D. H. Single-Cell Responses in Striate Cortex of Kittens Deprived of Vision in One Eye. J Neurophysiol 26, 1003–1017, doi:10.1152/jn.1963.26.6.1003 (1963).

63 Murphy-Baum, B. L. & Taylor, W. R. The Synaptic and Morphological Basis of Orientation Selectivity in a Polyaxonal Amacrine Cell of the Rabbit Retina. J Neurosci 35, 13336–13350, doi:10.1523/JNEUROSCI.1712-15.2015 (2015).

64 Vita, D. J., Orsi, F. S., Stanko, N. G., Clark, N. A. & Tiriac, A. Development and organization of the retinal orientation selectivity map. Nat Commun 15, 4829, doi:10.1038/s41467-024-49206-z (2024).

65 Garner, A. R. & Keller, G. B. A cortical circuit for audio-visual predictions. Nat Neurosci 25, 98–105, doi:10.1038/s41593-021-00974-7 (2022).

66 Keller, G. B., Bonhoeffer, T. & Hubener, M. Sensorimotor mismatch signals in primary visual cortex of the behaving mouse. Neuron 74, 809–815, doi:10.1016/j.neuron.2012.03.040 (2012).

67 El-Quessny, M., Maanum, K. & Feller, M. B. Visual Experience Influences Dendritic Orientation but Is Not Required for Asymmetric Wiring of the Retinal Direction Selective Circuit. Cell Rep 31, 107844, doi:10.1016/j.celrep.2020.107844 (2020).

68 Hamby, A. M., Rosa, J. M., Hsu, C. H. & Feller, M. B. CaV3.2 KO mice have altered retinal waves but normal direction selectivity. Vis Neurosci 32, E003, doi:10.1017/S0952523814000364 (2015).

69 Gollisch, T. & Meister, M. Eye smarter than scientists believed: neural computations in circuits of the retina. Neuron 65, 150–164, doi:10.1016/j.neuron.2009.12.009 (2010).

70 Sanes, J. R. & Masland, R. H. The types of retinal ganglion cells: current status and implications for neuronal classification. Annu Rev Neurosci 38, 221–246, doi:10.1146/annurev-neuro-071714-034120 (2015).

71 Happe, F. G. Studying weak central coherence at low levels: children with autism do not succumb to visual illusions. A research note. J Child Psychol Psychiatry 37, 873–877, doi:10.1111/j.1469-7610.1996.tb01483.x (1996).

72 Jones, B. W. et al. Retinal remodeling. Jpn J Ophthalmol 56, 289–306, doi:10.1007/s10384-012-0147-2 (2012).

73 Harpaz, R., Nguyen, M. N., Bahl, A. & Engert, F. Precise visuomotor transformations underlying collective behavior in larval zebrafish. Nat Commun 12, 6578, doi:10.1038/s41467-021-26748-0 (2021).

