## Supplemental Figure 1 for "Early visual experience elicits cellular and functional plasticity in the retina and alters behaviour"

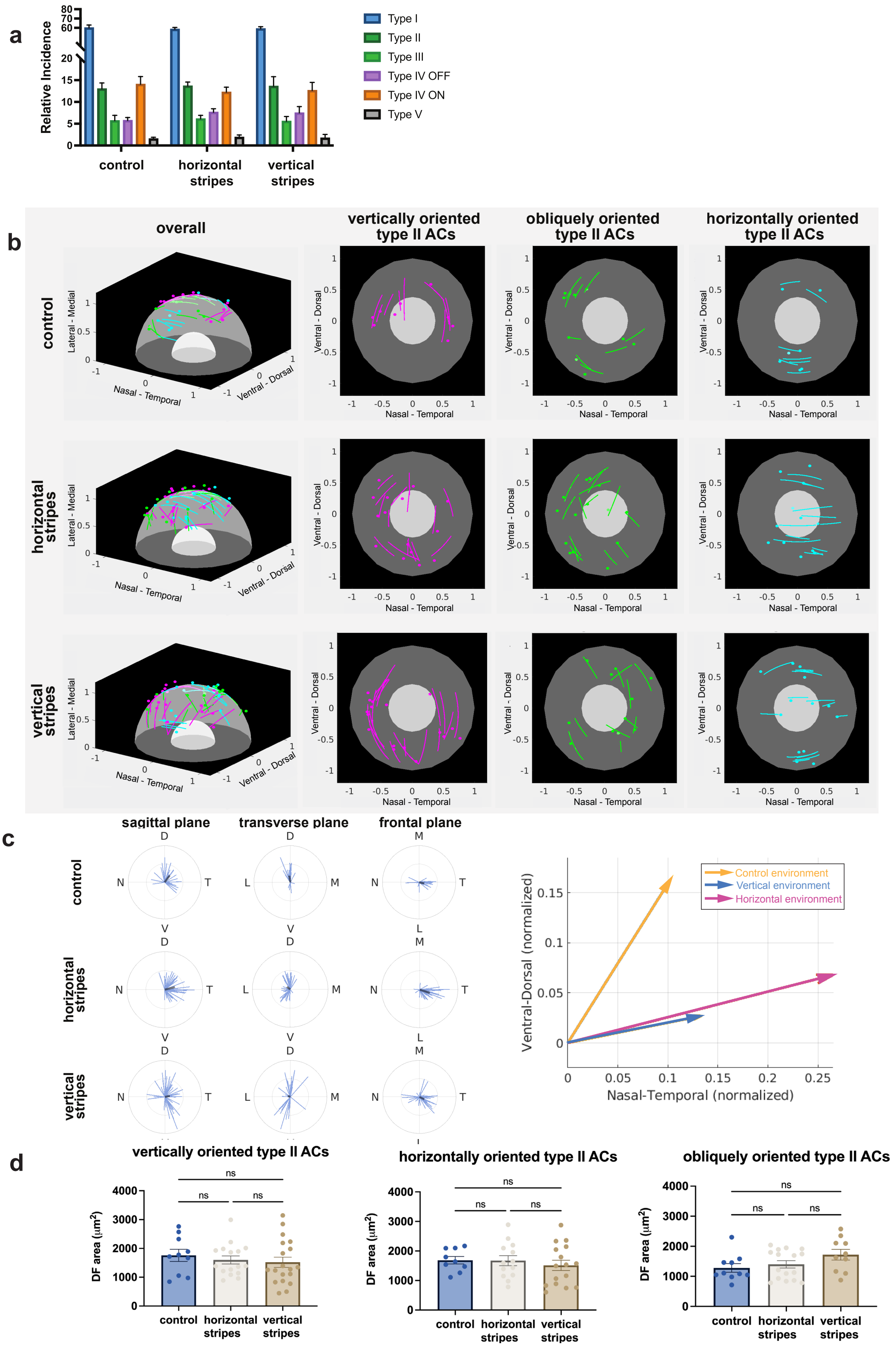


**Supplemental Figure 1: Exposure to different visual environments leads to changes in morphology of orientation-selective amacrine cells**

**a.** Relative incidence of the observed tenm3+ AC types of embryos raised from 72 hpf to 5 dpf in control, vertical striped, and horizontal striped environments (N = 675 cells from 300 embryos per condition). (AC types x Environments, *P = 0.9952*;

Environments, $P = 0.9524$: AC types, ****$P < 0.0001$; Repeats, > 0.9999). Two-way ANOVA followed by Tuckey multiple comparison test. **b.** Schematic illustration of cell bodies and dendritic fields registered in 3D. White and grey hemispheres represents the lens and IPL. Cells are color-coded according to their orientation observed manually. Magenta: vertical; green: oblique; cyan: horizontal. Lengths are normalized to the distance between the IPL to the centre of lens. **c.** (Left) Length and orientation of the dendritic fields projected in the three orthogonal planes. The averaged length and orientation are shown in black. (Right) The averaged length and orientation of cells in sagittal view. D, dorsal; IPL, inner plexiform layer; L, lateral; M, medial; N, nasal; T, temporal; V, ventral. **d.** Dendritic field areas of vertically, horizontally and obliquely oriented type II cells, split into the three environments the animals were raised in (i.e., control, horizontal stripes, and. vertical stripes). (Vertically oriented type II cells, $P = 0.6409$ (C vs. V); $P = 0.8331$ (C vs. H), $P = 0.9329$ (V vs. H); Horizontally oriented type II cells, $P = 0.7718$ (C vs. V); $P = 0.9983$ (C vs. H), $P = 0.7635$ (V vs. H); Obliquely oriented type II cells, $P = 0.1300$ (C vs. V); $P = 0.8197$ (C vs. H), $P = 0.2568$ (V vs. H); Graphs show mean and error bars ± s.e.m. One-way ANOVA followed by Tuckey multiple comparison test. C, control; V, vertical stripes, H, horizontal stripes.
