## Supplemental Figure 2 for "Early visual experience elicits cellular and functional plasticity in the retina and alters behaviour"

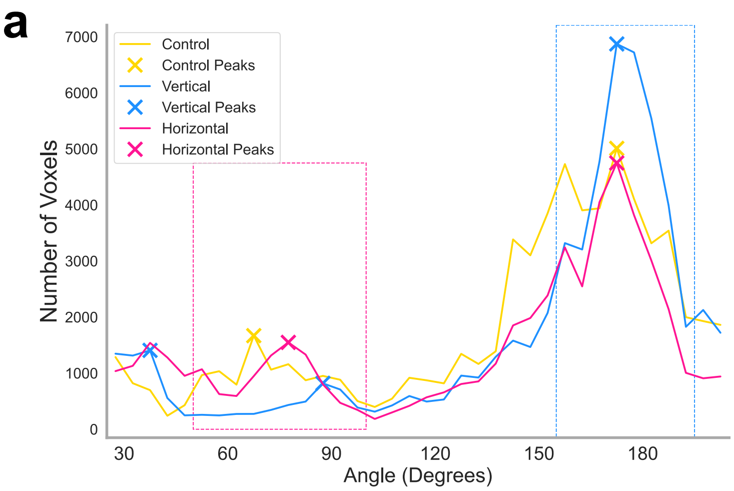


b

| Group | Population Vertical Index |
|---|---|
| C | 0.78 |
| V | 0.90 |
| H | 0.75 |

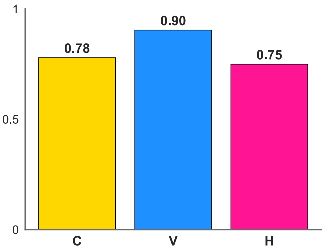


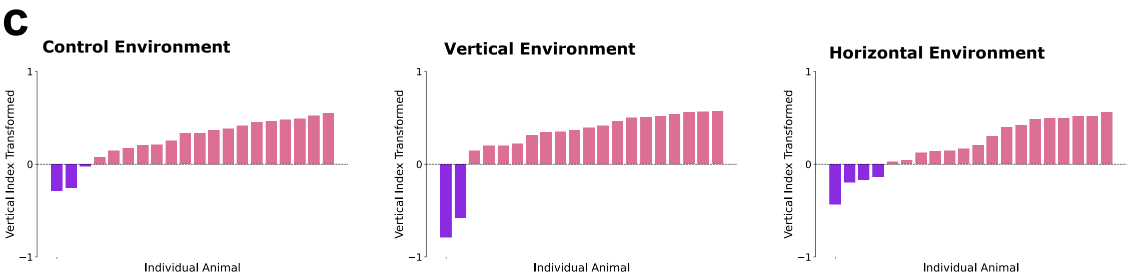


**Supplemental Figure 2: Relative changes of orientation-selective output from larvae raised in different conditions**

**a**. Plot showing best fit curve of OS responses across 0 to 180 degrees, representing all voxels with an OSI > 0.5 for larvae raised in control (yellow), vertical striped (blue), and horizontal striped (magenta) (n = 20 for each condition, data as in Figure 2). The vertical and horizontal visual stimuli peaks are marked with X. The dashed lines represent the set ranges for vertical or horizontal preference (between 0° to 15° and 150° to 180° for vertical preference, and 50° to 100° for horizontal preference). **b.** The table shows a vertical preference index for each environment calculated. Number of voxels is calculated as the area under the curve of the defined horizontal or vertical preference window. The bar graph is a visual representation of the vertical indices across environments. **c.** Vertical index values for each individual fish sorted according to their preference magnitude from horizontal (purple) to vertical (pink) .
