## Supplemental Figure 3 for "Early visual experience elicits cellular and functional plasticity in the retina and alters behaviour"

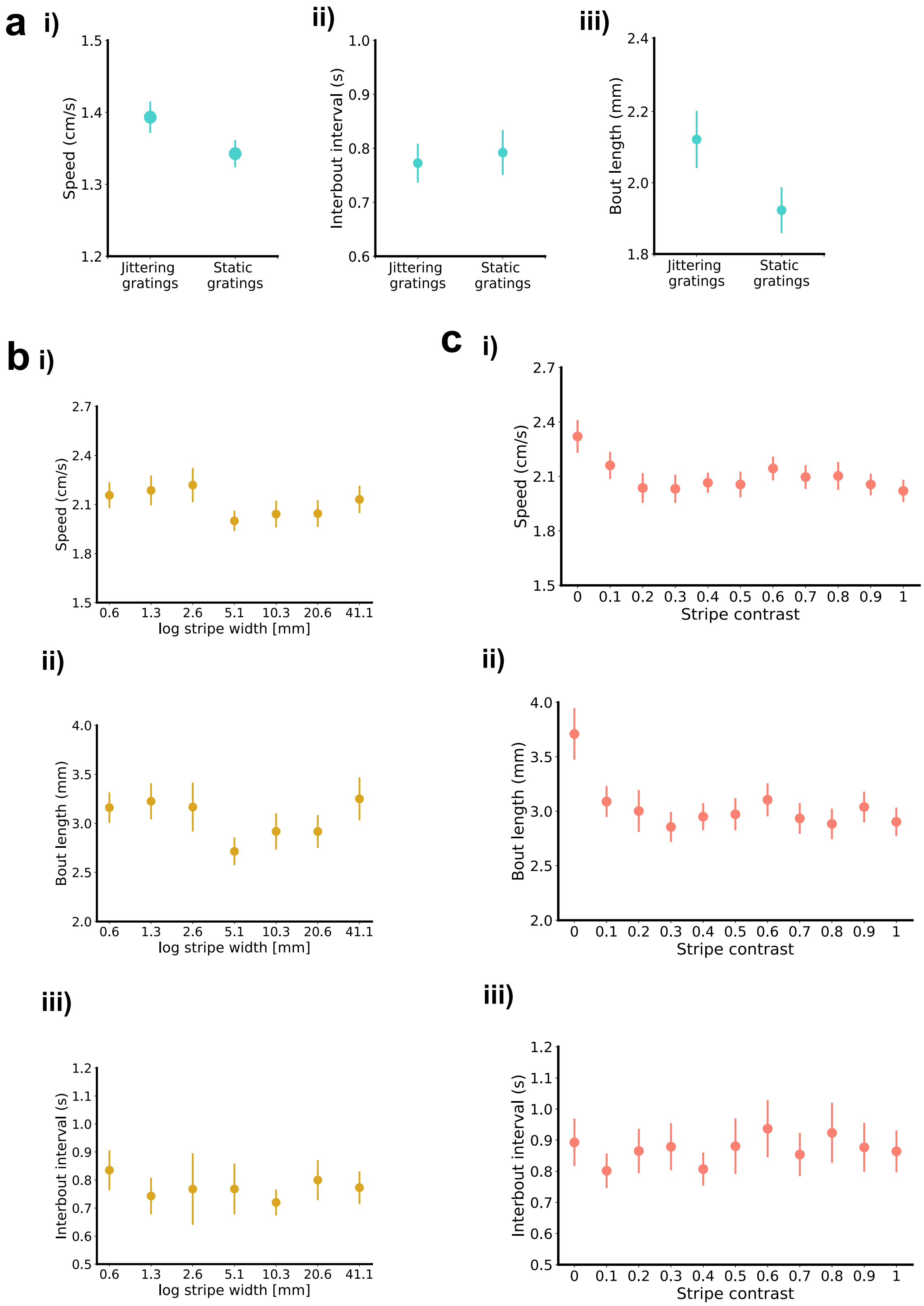


**Supplemental Figure 3: Speed, bout length and interbout interval in novel behavioural paradigm**

**a.** Jittering versus static: Average speed of bouts, interbout interval and bout length behaviours when presented with jittering and static stimuli. **b.** Stripe frequency: There was no difference in the speed, interbout interval or bout length of larvae when shown gratings of different widths. **c.** Stripe contrast: Speed considers the average speed of the larvae per stimuli, interbout interval measures the gap between each bout and length is the length per bout when shown different stimuli. **a-c.** All error bars are mean ± s.e.m. over fish. N numbers match values shown in Fig 4. P values are based on two-tailed t-test with Welch's correction. Asterisks indicate significance *(*P < 0.05, ****P < 0.0001).*
