## Supplemental Figure 4 for "Early visual experience elicits cellular and functional plasticity in the retina and alters behaviour"

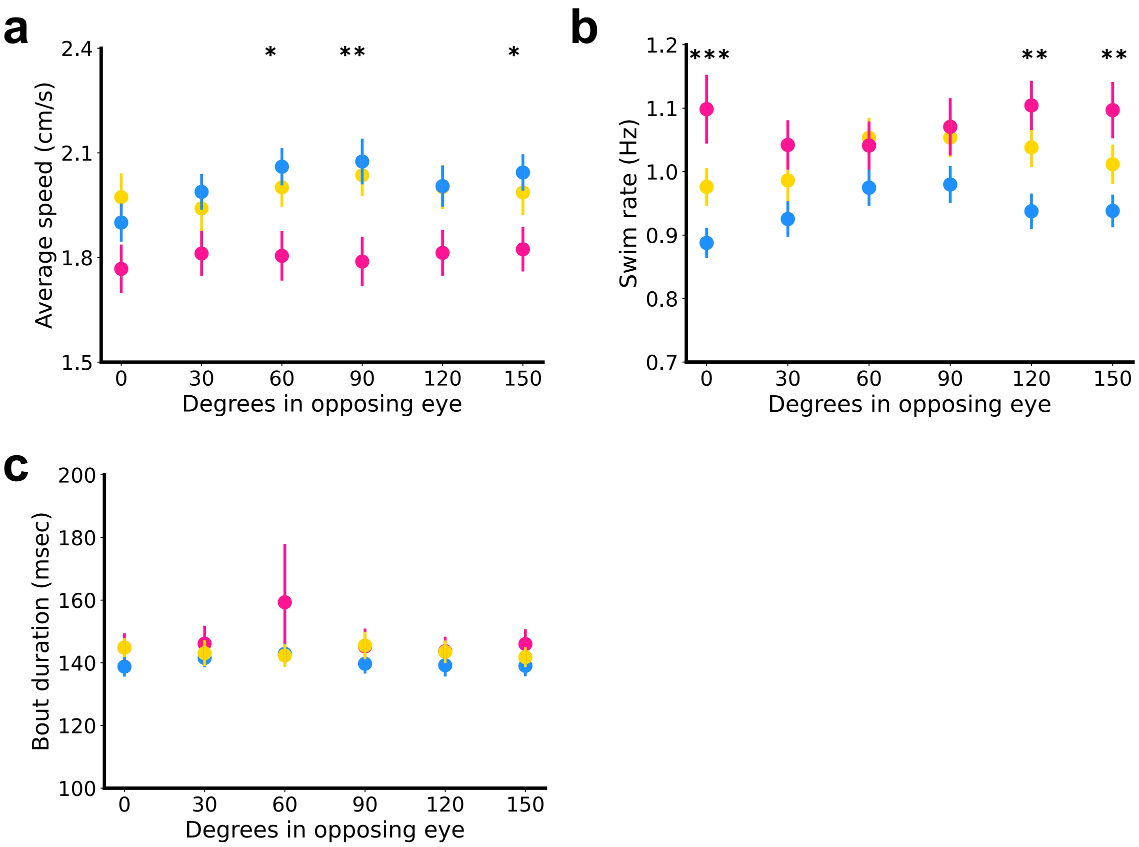


**Supplemental Figure 4: Behavioural responses in larvae raised in different environments**

**a.** Average speed of larvae (cm/s) per stimuli when shown parallel stimuli in one eye and stimuli of changing angles in the opposing eye (60 degrees, *$P = 0.0142$ (V vs. H); 90 degrees, *$P = 0.0137$ (C vs. H), **$P = 0.0048$ (V vs. H); 150 degrees, *$P = 0.0417$ (V vs. H). **b.** Frequency of bouts (Hz) per stimuli when shown parallel stimuli in one eye and stimuli of changing angles in the opposing eye (30 degrees, *$P = 0.0385$ (C vs. H), ***P = *0.0002* (V vs. H); 120 degrees, **$P = 0.0038$ (V vs. H); 150 degrees, **$P = 0.0064$ (V vs. H). **c.** Bout duration (msec) per stimuli when shown parallel stimuli in one eye and stimuli of changing angles in the opposing eye. All error bars are mean ± s.e.m. over fish. N numbers match values shown in Fig 5.
